# Negative regulation of Activation-Induced Cytidine Deaminase gene transcription in developing B cells by a PU.1-interacting intronic region

**DOI:** 10.1101/2024.05.29.596448

**Authors:** Allanna C. E. MacKenzie, Mia P. Sams, Jane Lin, Carolina Reyes Batista, Michelle Lim, Chanpreet K. Riarh, Rodney P. DeKoter

## Abstract

Activation-induced cytidine deaminase (AID, encoded by *Aicda*) plays a key role in somatic hypermutation and class switch recombination in germinal center B cells. However, off-target effects of AID are implicated in human leukemia and lymphoma. A mouse model of precursor B cell acute lymphoblastic leukemia driven by deletion of the related transcription factors PU.1 and Spi-B revealed C->T transition mutations compatible with being induced by AID. Therefore, we hypothesized that PU.1 negatively regulates *Aicda* during B cell development. *Aicda* mRNA transcript levels were increased in leukemia cells and preleukemic bone marrow pre-B cells lacking PU.1 and/or Spi-B, relative to wild type cells. Using chromatin immunoprecipitation, PU.1 was found to interact with a negative regulatory region (R2-1) within the first intron of *Aicda*. CRISPR-Cas9-induced mutagenesis of R2-1 in cultured pre-B cells resulted in upregulation of *Aicda* in response to lipopolysaccharide stimulation. Mutation of the PU.1 interaction site and neighboring sequences resulted in reduced repressive ability of R2-1 in transient transfection analysis followed by luciferase assays. These results show that a PU.1-interacting intronic region negatively regulates *Aicda* transcription in developing B cells.

## 1. Introduction

Precursor B-cell acute lymphoblastic leukemia (pre-B-ALL) is the most frequently occurring cancer in young children. Despite a high remission rate, pre-B-ALL is still a leading cause of cancer-related mortality in children (Inaba and Mullighan, 2020). Pre-B-ALL arises from precursor B cells that are proliferating in response to Interleukin-7 (IL-7) signaling (Clark et al., 2014). Cell division must be carefully segregated from immunoglobulin gene recombination during B cell development to avoid off-target mutations caused by the RAG1/2 recombinase machinery (Teng et al., 2015). Another potential source of mutagenesis during B cell development is Activation Induced Cytidine Deaminase (AID, encoded by *Aicda*). B cell leukemias frequently have mutational signatures indicating a developmental history of mutations induced by AID or RAG1/2 (Swaminathan et al., 2015; Thatikonda et al., 2023).

AID is a member of the APOBEC family of enzymes (Feng et al., 2020). The function of AID is to deaminate deoxycytidines (dC) within the WR<u>C</u>Y (W=A/T, R=A/G) motif into deoxyuridines (dU) during the processes of somatic hypermutation or class switch recombination in B cells (Feng et al., 2020). Mutations are generated by error-prone repair of dU into thymine, producing AID’s characteristic C->T transition mutation (Feng et al., 2021). AID is highly expressed in germinal center B cells where the processes of somatic hypermutation and class switch recombination occur during immune responses (Feng et al., 2020). However, AID has also been shown to be expressed during B cell development where it plays a role in establishing B cell tolerance (Cantaert et al., 2015; Kuraoka et al., 2017; Kuraoka and Kelsoe, 2011). Therefore, AID is tightly regulated in a cell type- and developmental stage-specific manner to avoid the generation of leukemia and lymphoma-promoting mutations.

Four genomic regions with activating or repressive function were identified to regulate *Aicda* gene transcription (Tran et al., 2010). Regions 1, 3 and 4 activate *Aicda* transcription in a cell type-specific manner. Region 2 is located in the first intron of *Aicda* and functions primarily as a negative regulator of transcription (Tran et al., 2010). Region 2 was determined to contain 14 distinct transcription factor binding motifs including sites for c-Myb, E2F, Pax-5, and E2A (Tran et al., 2010). However, a limitation of this study was that it used exclusively transient transfection analysis and reporter assays to study *Aicda* regulatory regions (Zan and Casali, 2013). Thus, a significant gap in our understanding of developmental stage-specific regulation of *Aicda* remains to be addressed.

Mice homozygous for a null allele of *Spib* encoding the E26-transformation-specific (ETS) transcription factor Spi-B, and with a B cell-specific deletion of *Spi1* encoding the related ETS transcription factor PU.1, under control of Mb1 (Cd79a)-Cre (Mb1-CreΔPB mice) have an arrest of B cell development at the small pre-B cell stage (Batista et al., 2017). Mb1-CreΔPB mice develop precursor B cell acute lymphoblastic leukemia (pre-B-ALL) at 100% incidence by a median 18 weeks of age (Batista et al., 2018; Lim et al., 2020). Whole-exome sequencing of Mb1-CreΔPB leukemias revealed two distinct mutational signatures: 1) C->A transversion mutations at low variant allele frequencies (VAF) within a trinucleotide context suggesting that they were caused by reactive oxygen species, and 2) C->T transition mutations at high VAF within a trinucleotide context suggesting that they were caused by AID (Lim et al., 2020). Based on these observations, we hypothesized that *Aicda* transcription is negatively regulated by PU.1 during B cell development.

To determine if *Aicda* is regulated by PU.1, we investigated *Aicda* mRNA transcript levels and found that they were increased in leukemia cells or preleukemic bone marrow pre-B cells lacking PU.1 and/or Spi-B. PU.1 interacted with a site (termed R2-1) within the negative regulatory Region 2 of the *Aicda* first intron. CRISPR-Cas9-induced mutagenesis of R2-1 resulted in upregulation of *Aicda* mRNA transcript levels in response to lipopolysaccharide-induced stimulation of the pre-B cell line 38B9. Transient transfection and luciferase reporter analysis of R2-1 suggested that transcription factors in addition to PU.1 are involved in negative regulation of *Aicda*. These results show that PU.1 is involved in the constraint of *Aicda* transcription during B cell development by interaction with an intronic regulatory region.

## 2. Materials and methods

### 2.1. Cell Culture

38B9 pre-B cells (Alt et al., 1982) were cultured at 37°C and 5% CO_2_ atmosphere in RPMI-1640 media (Wisent, Saint-Jean-Baptiste, QC) containing 10% fetal bovine serum (Wisent), supplemented with 1X penicillin/streptomycin/L-glutamine (Wisent), and 5×10^−5^ M β-mercaptoethanol (Millipore-Sigma, Oakville ON). 38B9 cells were stimulated in culture with 10 μg/ml lipopolysaccharide from *E. coli* 0111:B4 (Millipore-Sigma). Wild type pro-B and 973 PU.1/Spi-B-deficient pro-B cells (Lim et al., 2020) were cultured in Iscove’s Modified Dulbecco’s Media (Wisent) with 10% fetal bovine serum, supplemented with 1x penicillin/streptomycin/L-glutamine, 5×10^−5^ M β-mercaptoethanol, and supplemented with interleukin-7 (IL-7)-conditioned media from the J558-IL-7 cell line (Batista et al., 2017).

### 2.2. Flow Cytometry and Cell Sorting

Bone marrow cells were prepared from C57Bl/6J mice and red blood cells were removed by hypotonic lysis. Cells were stained with antibodies including Brilliant Violet 421-anti-B220 (RA3-6B2, Biolegend, San Diego CA), fluorescein isothiocyanate-anti-BP-1 (6C3, Biolegend), allophycocyanin-anti-IgM (II/41, BD Biosciences, Mississauga, ON), biotin anti-CD43 (S7, BD Biosciences), and Phycoerythrin/Cy5 streptavidin (Biolegend). Bone marrow cell and 38B9 cell line sorting was performed using a FACSAria III cell sorter (BD Immunocytometry systems, San Jose, CA).

### 2.3. Molecular Cloning and DNA Sequence Analysis

Modifications to the pGL3-promoter plasmid (Promega, Madison, WT) were made using restriction enzymes and T4 DNA ligase from New England Biolabs (NEB, Whitby, ON). The immunoglobulin heavy chain enhancer (EiH) was PCR-amplified and ligated into the BamHI site, and *Aicda* R2-1 was PCR amplified and ligated into the KpnI site to generate pGL3-promoter-EiH-R2.1. PCR products were cloned using the TA-Blunt-Zero cloning kit (GeneBio Systems Inc., Burlington, ON). Further modifications were performed by site-directed mutagenesis using the Q5 Site-Directed Mutagenesis Kit (NEB). All modifications to the 2X_pX458_pSpCas9(BB)-2A-GFP_SOX17_bKO plasmid (Addgene, Watertown, MA) were generated by Q5 site-directed mutagenesis. Primers were designed using the NEBaseChanger tool (NEB) and are shown in **Supplemental Table 1**. Sanger DNA sequencing was used to confirm all constructs. For all DNA sequence analyses, mouse Mm10 genome was used as the reference sequence. ENCODE candidate cis-regulatory elements were based on DNase 1 hypersensitivity assays as described (Davis et al., 2018). Transcription factor binding sites were predicted using MATCH (gene-regulation.com) or PROMO (Messeguer et al., 2002). DNA sequence comparisons were performed using MacVector 18.6.4 (MacVector, Apex, NC).

### 2.4. Chromatin Immunoprecipitation

1 × 10^7^ 38B9 cells were crosslinked in 1% formaldehyde for 10 minutes. Crosslinking was stopped by addition of 0.125 M glycine (Millipore-Sigma). Crosslinked cells were washed with cold PBS before pellets were flash-frozen in liquid nitrogen. Chromatin was sonicated using a Biorupter-300 (Diagenode, Denville, NJ) at high power for 30 cycles. Chromatin solution containing 300 µg DNA was immunoprecipitated using Protein G Dynabeads (Thermo-Fisher Scientific, Mississauga, ON) conjugated to rabbit polyclonal anti-PU.1 antibody (SC-352, Santa Cruz Biotechnology, Santa Cruz, CA). Immunoprecipitated chromatin was washed and eluted before being de-crosslinked at 65°C overnight. DNA was isolated using a Zymo DNA purification kit (Zymo Research, Irvine, CA). PU.1 interaction was determined by qPCR and calculated as per cent enrichment relative to input chromatin using primers recognizing region R2-1 and a downstream negative control region. Primer sequences are shown in **Supplemental Table 1**.

### 2.5. Quantitative Polymerase Chain Reaction

Total RNA was extracted from cultured or sorted cells using the RNeasy kit (Qiagen, Toronto, ON). Complementary DNA was synthesized using the iScript cDNA Synthesis kit (Bio-Rad, Mississauga, ON). RT-qPCR or qPCR was performed with Sensifast SYBR Green No-Rox master mix (Meridian Biosciences, Memphis TN) using the QuantStudio 5 PCR System (Thermo Fisher Scientific). Gene expression was measured in triplicate for all biological replicate experiments using the ΔΔCT method. Primer sequences are shown in **Supplemental Table 1.**

### 2.6. Transient Transfection and Luciferase Assay

Luciferase assays were performed using the dual luciferase Reporter Assay System (Promega). The pGL3-promoter-EiH plasmid served as positive control, while pGL3-basic and pGL3-promoter-EiH-R1 were negative controls in different experiments (Promega). For each transfection, 4×10^6^ 38B9 cells were transfected with 0.5 µg of the pRL-TK plasmid encoding Renilla luciferase and 10 µg of pGL3-based plasmid encoding firefly luciferase. Electroporation was performed using 4 mm gap cuvettes (Fisher Scientific Canada, Toronto, ON) and a Gene Pulser II instrument (Bio-Rad) with settings of 950 µF and 220V. Following incubation for 48hr, transfected cells were lysed with passive lysis buffer and subjected to 3 rounds of freeze-thaw. Dual luciferase activity was measured using a BioTek Cytation 5 Cell Imaging Multimode Reader (Agilent, Santa Clara, CA). Normalized luciferase activity was determined by calculating Firefly to Renilla signal ratio.

### 2.7. Statistical analysis

Statistical analyses were determined using Prism 8.4.3 (GraphPad, San Diego, CA) and were based on independent biological replicate experiments. Statistical tests and group sizes are indicated in the figure legends.

## 3. Results

### 3.1. Increased levels of Aicda expression in PU.1/Spi-B deficient B cells

To determine if transcription of the *Aicda* gene encoding AID is dysregulated by genetic deletion of PU.1 and/or Spi-B in developing B cells, relative mRNA transcript levels of *Aicda* were compared between cultured wild type pro-B cells and PU.1/Spi-B knockout leukemia cells (973 cells) from Mb1-CreΔPB mice (Lim et al., 2020). There was a 2.3-fold higher level of *Aicda* mRNA transcripts in 973 cells compared to wild type pro-B cells (**Fig. 1A**). To determine if *Aicda* is affected by PU.1 and Spi-B in preleukemic developing B cells, “Hardy” fractions A-D were enriched from the bone marrow of wild type (C57Bl/6), Spi-B knockout (ΔB), or PU.1/Spi-B double knockout (Mb1-Cre1′PB) mice by cell sorting (Hardy et al., 1991). Fraction A cells were B220+ CD43+ BP1-HSA-, fraction B B220+ CD43+ BP1-, HSA+, fraction C B220+ CD43+, BP1+ HSA+, and fraction D B220+ CD43-IgM-. RT-qPCR analysis showed that *Aicda* was elevated in fraction B pro-B cells enriched from PU.1/Spi-B double knockout mice relative to Spi-B knockout or WT mice (**Fig. 1B**). These results suggested that PU.1 negatively regulates *Aicda* transcription in B cells.

**Figure 1.**
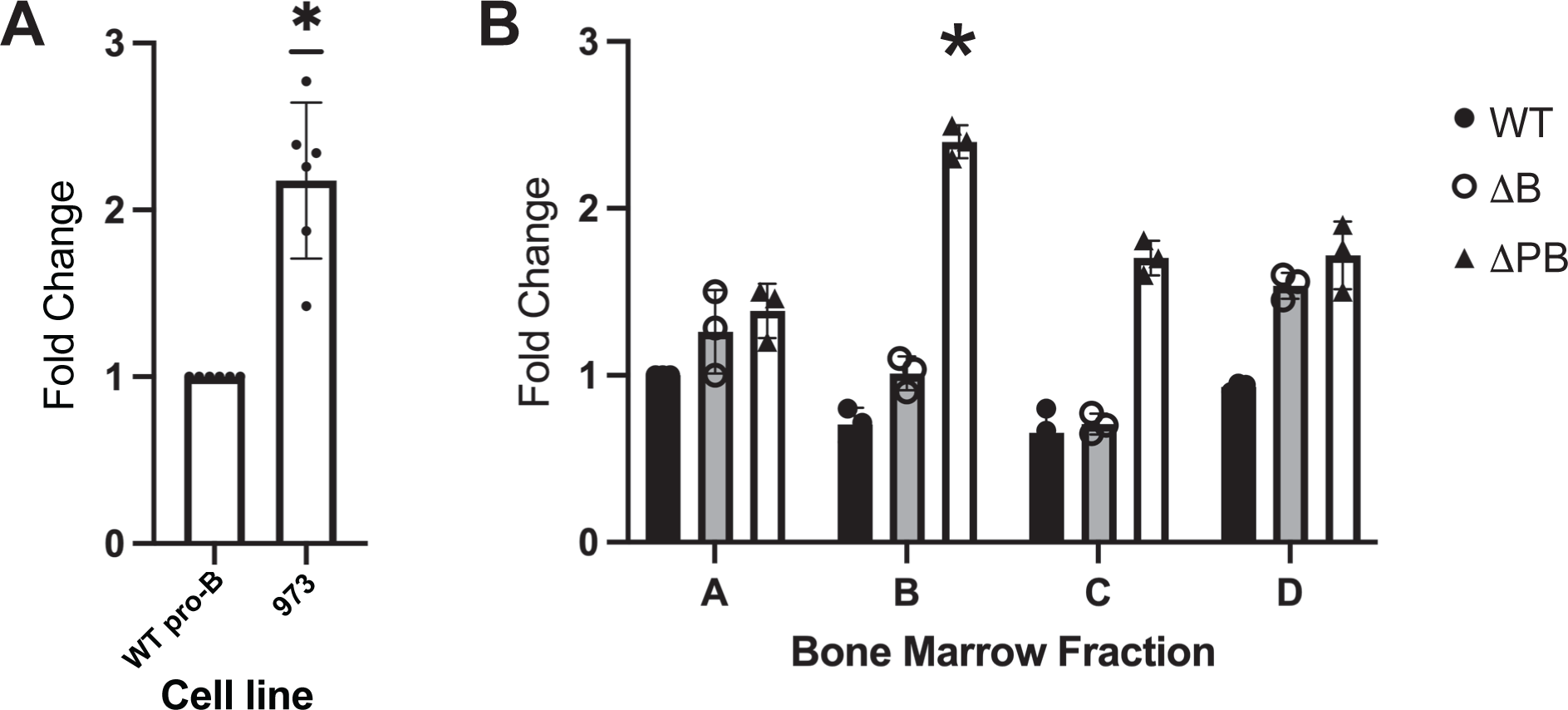
Increased mRNA transcript levels of *Aicda* in cells lacking PU.1 and Spi-B. A) increased mRNA transcript levels of *Aicda* in cultured 973 leukemia cells lacking PU.1 and Spi-B relative to wild type fetal liver-derived pro-B cells. *p=0.031 by one-sample t and Wilcoxon test. B) Increased mRNA levels of *Aicda* in fraction B bone marrow pro-B cells from mice lacking PU.1 and Spi-B (1′PB) relative to wild type (WT) or mice lacking Spi-B only (1′B). *p=0.013 by Kruskal-Wallis and Dunn’s Multiple comparisons test.

### 3.2. Identification of PU.1 and IKZF3 sites in a negative regulatory region of Aicda

Region 2 within the first intron of *Aicda* was reported to function as a negative regulator of transcription (Tran et al., 2010) (**Fig. 2A**). To determine if PU.1 interacts with *Aicda* regulatory regions, anti-PU.1 ChIP-seq data previously generated by our lab was analyzed (Batista et al., 2017). Four peaks of PU.1 interaction were identified that were located at ENCODE candidate cis-regulatory elements within the *Aicda* locus (**Fig. 2A**). Two PU.1 peaks were located within *Aicda* region 2 and were named R2-1 and R2-2 (**Fig. 2A**). ENCODE ChIP-seq data showed that PU.1 interacted with R2-1 in human cells (data not shown). Both R2-1 and R2-2 showed a high degree of multispecies DNA sequence identity based on placental conservation and a 60-species multispecies conservation analysis (MultiZ conservation) (**Fig. 2A**). PU.1 and IKZF1/3 sites have previously been shown to share a common GGAA core motif and can interact with a common set of binding sites in the genome (Katerndahl et al., 2017; Zarnegar and Rothenberg, 2012). To determine if R2-1 can interact with IKZF family members, we re-analyzed anti-IKZF3 ChIP-seq data from our lab (Rodrigues de Oliveira et al., 2024). The result showed that IKZF3 interacts specifically with R2-1 and not other ENCODE predicted cis-regulatory elements (**Fig. 2A**).

**Figure 2.**
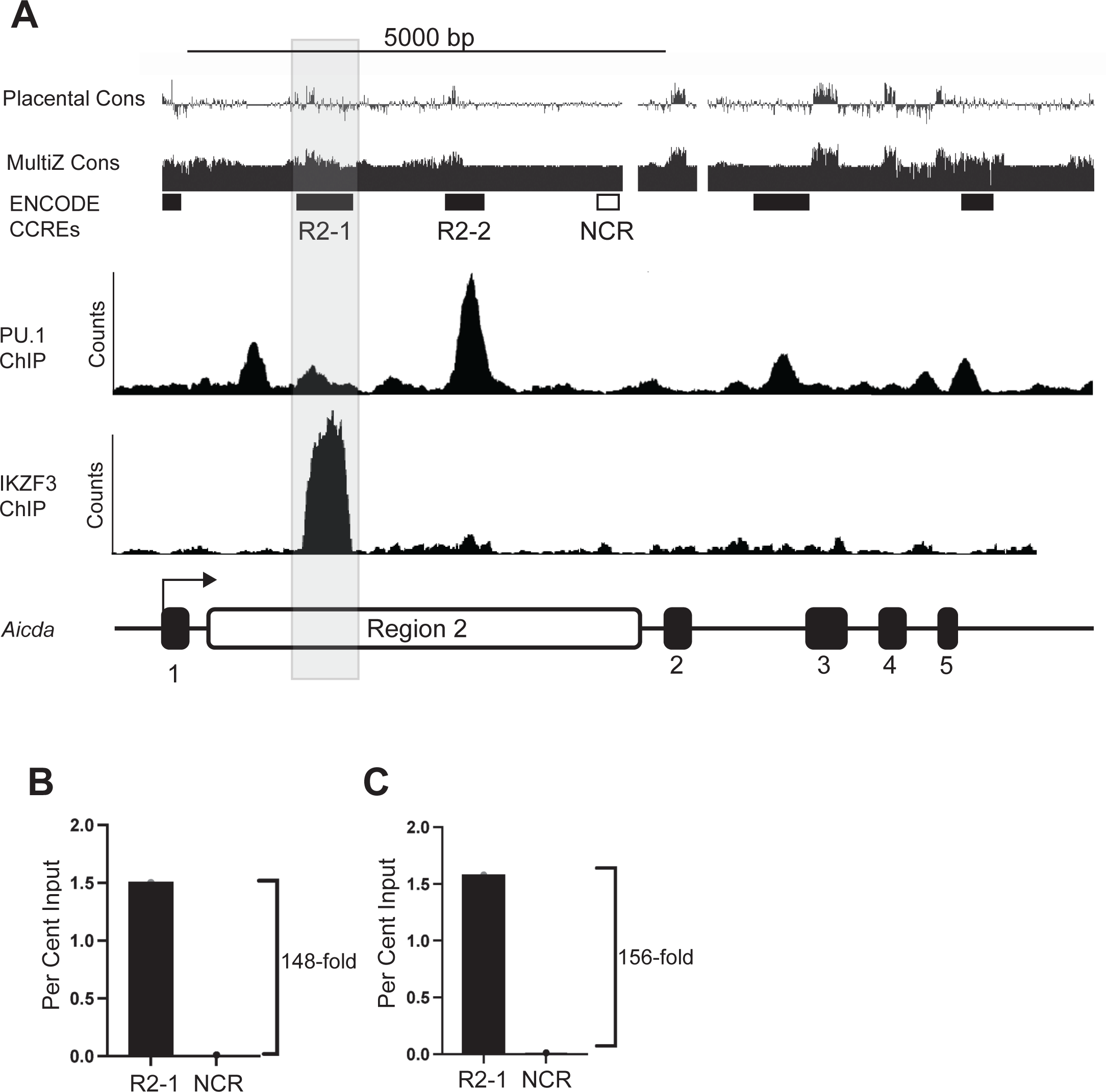
Identification of a PU.1 interaction site within a negative regulatory region of *Aicda*. A) Interaction of PU.1 and IKZF3 with sites in the *Aicda* gene. Top row shows shows placental mammal nucleotide conservation analysis from UCSC Genome Browser (Placental Cons). Second row shows nucleotide conservation analysis of 60 species from UCSC Genome Browser (MultiZ Cons). Third row shows ENCODE candidate cis-regulatory elements from UCSC Genome Browser (ENCODE CCREs). Regions 2-1 and 2-2 are indicated as R2-1 and R2-2. Negative control region (NCR) is indicated by an open box. Gray shading indicates region 2-1. Fourth row shows anti-PU.1 ChIP-seq analysis (PU.1 ChIP). Fifth row shows anti-IKZF3 ChIP-seq analysis (IKZF3 ChIP). Bottom row shows schematic of the Aicda gene, with exons shown in block, region 2 shown in white, and the transcription start site indicated by an arrowheard. B, C) Association of PU.1 with R2-1 as indicated by anti-PU.1 chromatin immunoprecipitation analysis. Y-axis indicates per cent input; the x-axis indicates the primer set used, either R2-1 or negative control region (NCR). B and C represent two independent biological replicate experiments.

To confirm that PU.1 interacts with R2-1 in mouse cells, anti-PU.1 ChIP was performed in 38B9 leukemia cells followed by qPCR analysis using primers recognizing R2-1 or a negative control region (NCR) (**Fig. 2A**). R2-1 was 148-fold or 156-fold enriched by anti-PU.1 ChIP compared to the NCR in two independent experiments (**Fig. 2B, 2C**). In conclusion, PU.1 interacts with region R2-1 of *Aicda* in mouse 38B9 cells. Therefore, R2-1 was prioritized as a candidate PU.1-regulated regulatory element.

### 3.3. Region R2-1 has transcriptional repression activity in transient transfection analysis

Examination of the ∼300 bp R2-1 and R2-2 DNA sequences using JASPAR CORE identified one predicted PU.1 binding site in each region that was located at the centre of the PU.1 ChIP-seq peak. PCR was used to amplify ∼500 bp R2-1 and R2-2 fragments encompassing the identified PU.1 binding site (**Fig. 3A**). To determine if R2-1 or R2-2 function as a repressive transcriptional element, these DNA segments were ligated into luciferase reporter plasmids. R2-2 had no activity as either an enhancer or an activator (data not shown). However, experiments performed with R2-1 cloned into the pGL3-promoter vector suggested that R2-1 functioned as a transcriptional repressor. To test transcriptional repression from a promoter-enhancer combination highly active in B cells, the pGL3-promoter luciferase reporter vector was modified to insert the immunoglobulin heavy chain (*IgH*) intronic enhancer (EiH) (Banerji et al., 1983) in the downstream position (**Fig. 3B**). Electroporation of 38B9 mouse leukemia cells followed by dual luciferase assays showed that inclusion of the EiH enhancer increased luciferase activity of the pGL3-promoter vector by more than 10-fold (**Fig. 3C**). Next, the R2-1 DNA fragment was ligated upstream of the minimal SV40 promoter in the pGL3-promoter-EiH vector (**Fig. 3B**). Electroporation followed by dual luciferase assays showed that the R2-1 element reduced luciferase activity by 2.2-fold (**Fig. 3D**). This result supported R2-1 functioning as a transcriptional repressive element, in agreement with the report of region 2 as a transcriptional silencer element (Tran et al., 2010).

**Figure 3.**
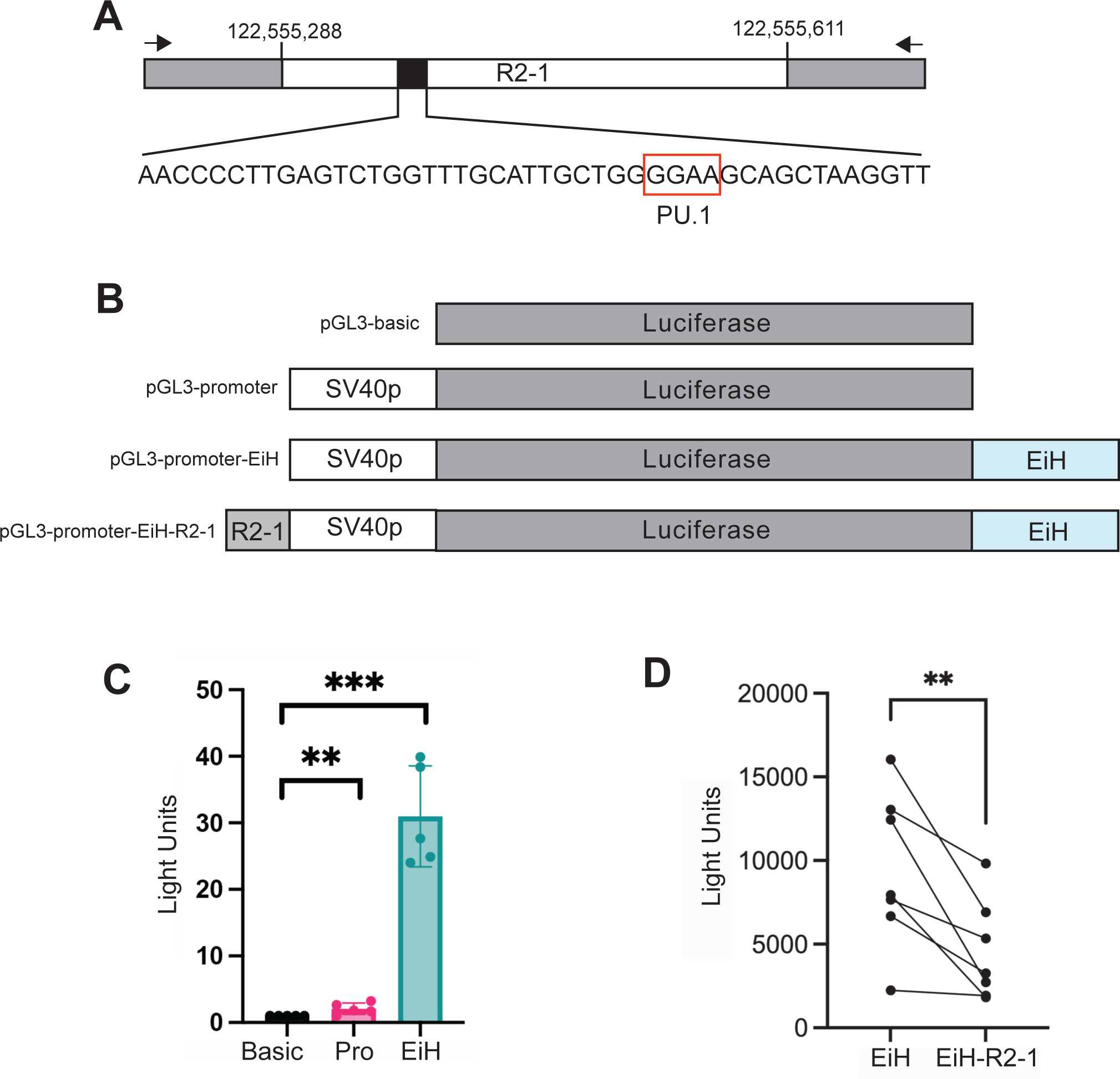
R2-1 of *Aicda* has repressive activity in luciferase reporter assays. A) Schematic of region 2-1 (R2-1) in the *Aicda* gene locus. Filled black box indicates the location of the PU.1 binding site. White indicates the R2-1 region. Gray indicates additional “arms” added by PCR amplification using primers indicated by arrowheads. Numbers indicate Mm10 genome coordinates. The purine-rich core motif of PU.1 (GGAA) is indicated by a box. B) Schematic of pGL3-based luciferase reporter vectors. SV40p indicates SV40 minimal promoter. EiH indicates the Immunoglobulin Heavy Chain intronic enhancer. R2-1 indicates *Aicda* R2-1. C) The IgH intronic enhancer increases transcriptional activity of the SV40 minimal promoter. Dual luciferase assay was used to measure normalized light units (y-axis) following electroporation of 38B9 pre-B cells with pGL3-basic (Basic), pGL3-promoter (Pro), or pGL3-promoter-EiH (EiH). **p<0.01, ***p<0.001 by one-way ANOVA. D) R2-1 shows repressive activity. Dual luciferase assay was used to compare activity of pGL3-promoter-EiH (EiH) with pGL3-promoter-EIH-R2-1 (EiH-R2-1). **p=0.0053 by paired t-test. Paired biological replicate experiments are connected by lines.

### 3.4. Targeting the R2-1 PU.1 binding site using CRISPR-Cas9 mutagenesis

To determine if the R2-1 PU.1-binding site has transcriptional repression function *in vivo*, we set out to introduce mutations in 38B9 cells using CRISPR-Cas9. The plasmid pX458 (Wu et al., 2021), was selected because of its all-in-one features including a U6 promoter driving two sgRNA cassettes, CMV promoter, and 2A-EGFP as a selectable marker (**Fig. 4A**). pX458 was modified to remove one of the two sgRNA cassettes, followed by site-directed mutagenesis to insert a crRNA sequence targeting the GGAA (PU.1) site using an adjacent PAM sequence (**Fig. 4B**). After the final plasmid was confirmed by nanopore sequencing, it was transfected by Neon electroporation into 38B9 cells. 48 hours after transfection, single GFP-positive cells were enriched by cell sorting into 96-well plates containing complete media (**Fig. 4C**). Of 384 cells sorted, 46 clones grew for a cloning efficiency of 11%. Genomic DNA was isolated from each clone and the R1 region was PCR amplified followed by Sanger sequencing analysis. **Fig. 4B** shows the distribution of CRISPR-Cas9-induced mutations in 44 clones that were successfully sequenced. 5 clones had no detectable mutations, and 12 clones had single point mutations to the fourth nucleotide within the GGAA motif. Four clones had a 5 bp deletion adjacent to the PAM sequence. All other clones had unique patterns of point mutations, small deletions, and larger deletions overlapping the target GGAA sequence (**Fig. 4B**).

**Figure 4.**
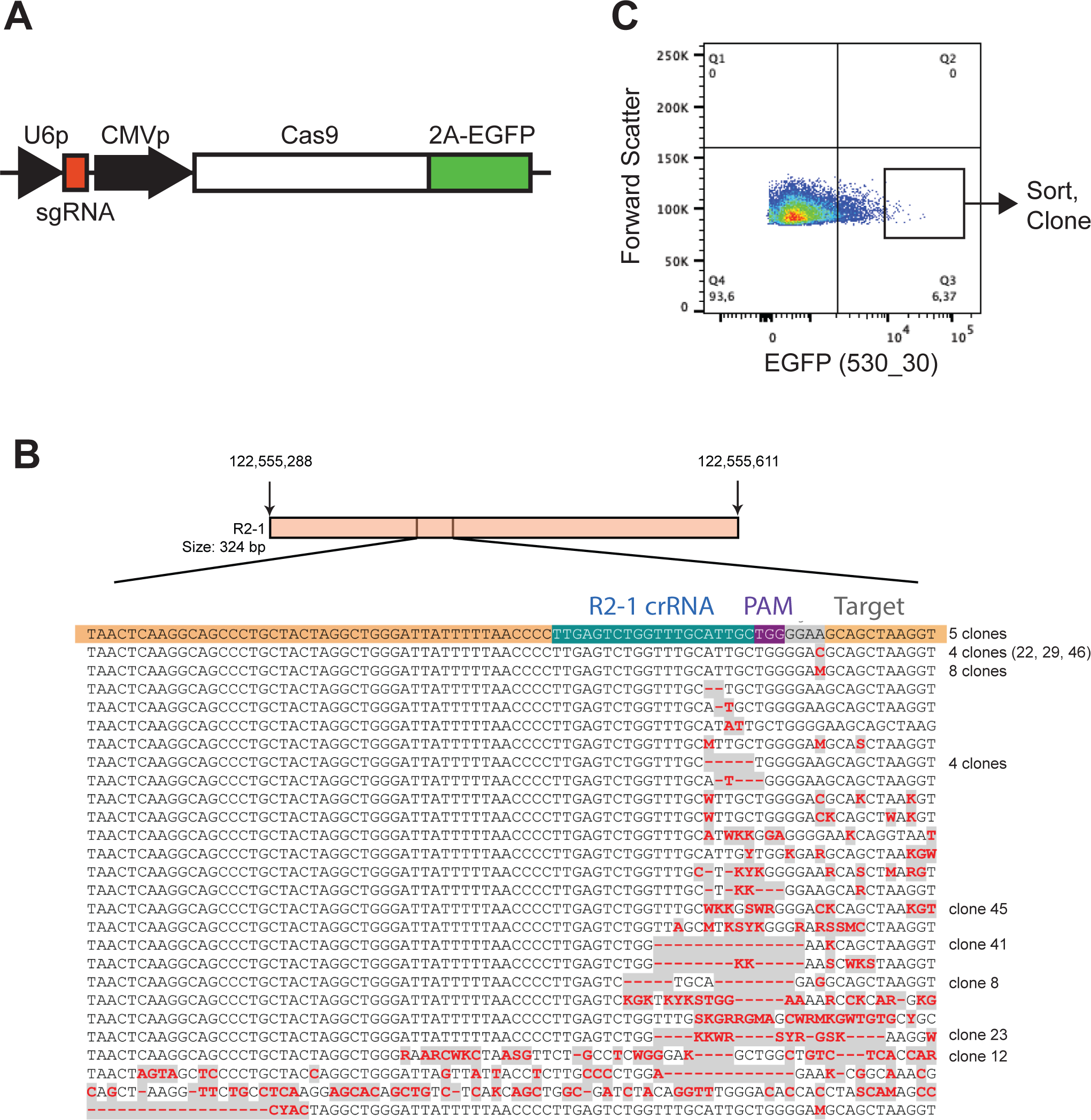
CRISPR-Cas9 mutagenesis of *Aicda* R2-1 in 38B9 pre-B cells. A) Schematic of the all-in-one CRISPR plasmid used, including U6 promoter (U6p), single guide RNA (sgRNA), cytomegalovirus promoter (CMVp), Streptococcus pyogenes Cas9 (Cas9), and 2A-enhanced green fluorescent protein (2A-EGFP). B) DNA sequences of R2-1 mutant clones. Top schematic shows location of sgRNA within the R2-1 sequence, including Crispr RNA (crRNA) in green, PAM sequence in purple, PU.1 GGAA core motif in gray, and target sequence in orange. DNA sequences and the number of 38B9 clones obtained for each sequence is shown in the bottom panel. C) Gating strategy for sorting of electroporated 38B9 cells. Y-axis indicates forward scatter, x-axis indicates green fluorescence (EGFP 530_30), and box indicates single-cell sort gate.

CRISPR-Cas9 mutagenesis typically creates double-stranded DNA breaks that are repaired by non-homologous end joining, generating a pool of heterozygous insertion or deletion mutations (Miyaoka et al., 2016; Paquet et al., 2016). However, in some cases, homology-directed repair can precisely fix these DNA breaks resulting in homozygous mutations (Paquet et al., 2016). To determine if the mutations made by the R2-1 CRISPR plasmid were heterozygous or homozygous, the R2-1 regions were PCR-amplified, the PCR products were cloned using a topoisomerase PCR cloning kit, and individual clones were sequenced. Five R2-1 clones amplified from clone 46 were sequenced to determine the frequency of the GGAA -> GGAC point mutation. Three clones had the GGAC point mutation, while two clones had the wild type GGAA sequence, suggesting that this mutation was heterozygous (**Supplemental Fig. 1A)**. For clone 8, 7 clones had the double deletion plus point mutation sequence, and 7 clones had a two-nucleotide deletion mutation (**Supplemental Fig. 1B**). Therefore clone 8 was heterozygous for two different short deletion mutations induced by CRISPR-Cas9.

### 3.5. Dysregulation of Aicda expression by R2-1 CRISPR-Cas9 mutagenesis

Lipopolysaccharide (LPS) is a potent stimulus of isotype switching in B cells and can initiate a proinflammatory response that activates *Aicda* (Park et al., 2005). We found that 10 μg/ml LPS robustly stimulated *Aicda* mRNA transcript levels in 38B9 cells (**Fig. 5A**). Three clones with point mutations in the PU.1 binding site (GGAA -> GGAC) were tested for *Aicda* induction in response to LPS. There were no significant differences in *Aicda* induction for clones 22, 29, and 46 compared to the unmutated parental 38B9 line following LPS stimulation (**Fig. 5B**). These results suggested that heterozygous mutation of the GGAA motif alone did not influence *Aicda* mRNA transcript levels *in vivo*.

**Figure 5.**
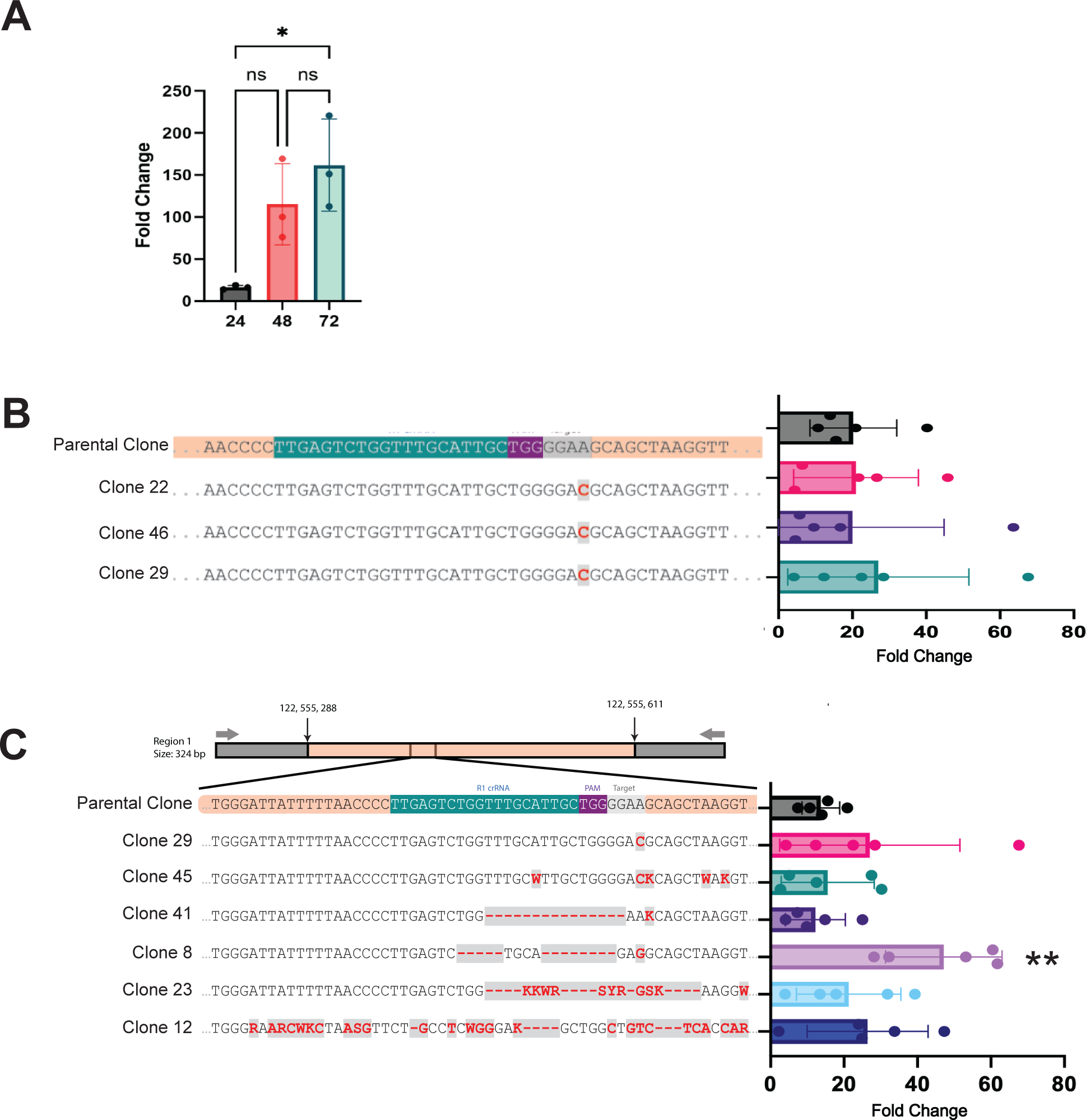
Dysregulation of *Aicda* transcription in 38B9 cells with a heterozygous R2-1 mutation. A) Stimulation of *Aicda* mRNA transcript levels by lipopolysaccharide treatment of 38B9 cells. Y-axis indicates fold change, x-axis indicates hours of treatment. Ns, not significant. *p=0.0037 by one sample t and Wilcoxon test. B) LPS Stimulation of *Aicda* mRNA transcript levels in clones with heterozygous point mutations of the PU.1 binding site. DNA sequences of clones 22, 46, and 29 are indicated on the left. Fold change in *Aicda* mRNA transcript levels from five independent biological replicate experiments are shown on the right. C) LPS Stimulation of *Aicda* mRNA transcript levels in clones with various heterozygous mutations. DNA sequences of clones are indicated on the left. Fold change in *Aicda* mRNA transcript levels from five independent biological replicate experiments are shown on the right. Clone 8 was **p=0.0076 by one-way ANOVA.

Next, clones with various point mutations or short deletions were tested. Clones 41 and 45 had small but not significant decreases in *Aicda* transcript level induction. Clones 23 and 12 had small but not significant increases in *Aicda* transcript level induction. Finally, clone 8 had a significant increase in induction of *Aicda* transcript levels following LPS stimulation compared to any other clone (**Fig. 5C**). This result suggested that heterozygous mutation of R2-1 in clone 8, as described above, resulted in the dysregulation of *Aicda* transcription.

### 3.6. Potential transcription factor binding locations within R2-1

To determine what predicted transcription factor binding may be disrupted by the clone 8 mutation, a 64 bp DNA sequence encompassing the GGAA motif was analyzed using MATCH or PROMO transcription factor prediction software. Using MATCH, clone 8 mutations were predicted to disrupt POU (Oct-1), ETS (Elk-1), IKZF1, and c-Rel binding sites (**Fig. 6A**). Using PROMO, clone 8 mutations were predicted to disrupt POU (Pou2f1), c/EBPΔ, XBP-1, FoxP3, ETS (Elk-1, Ets-1), STAT, and c/EBPα sites (**Fig. 6A**). Both prediction algorithms agreed that ETS binding sites and POU binding sites were disrupted. The prediction by MATCH that an IKZF1 site overlaps with the PU.1 binding site was intriguing because of our observation that R2-1 interacts with IKZF3 (**Fig. 2A**). In summary, these data suggest that ETS, POU, and IKZF1/3 binding sites may be disrupted by mutations found in clone 8.

**Figure 6.**
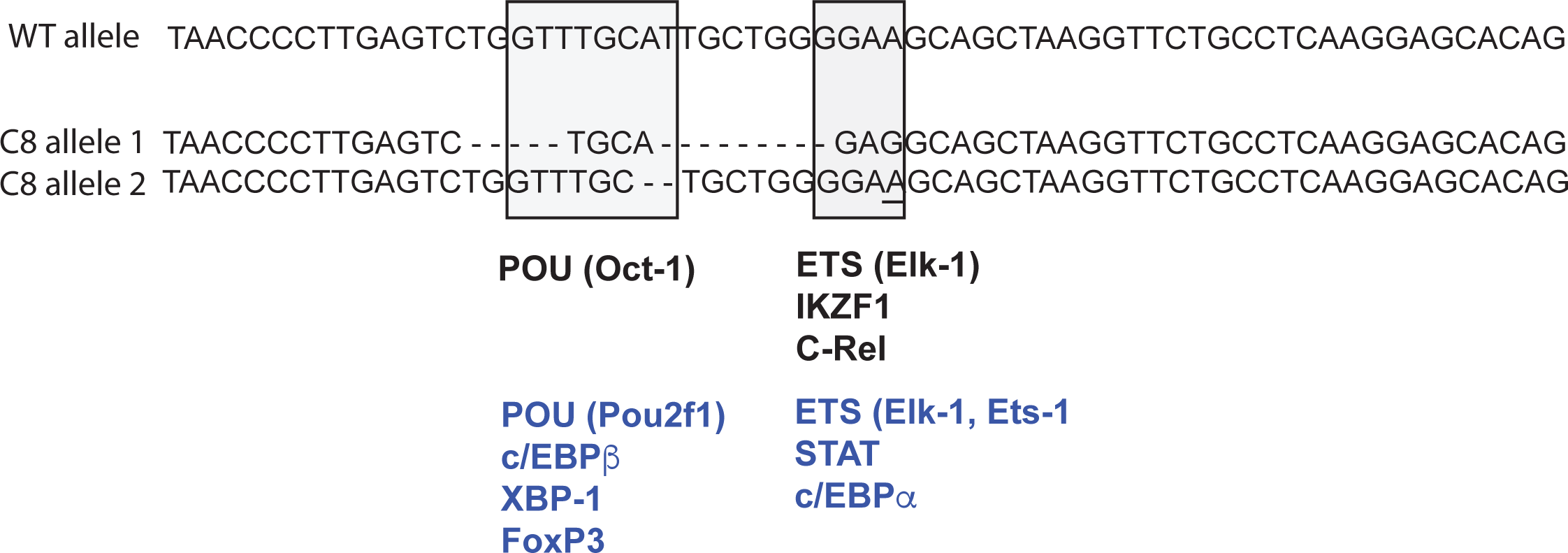
Prediction of disrupted transcription factor binding sites in mutated *Aicda* R2-1. DNA sequences of the wild type R2-1 allele (WT, top row) and clone 8 mutant alleles (C8 allele 1 and allele 2, bottom row) are shown with the POU core CTTTGCAT and PU.1 core GGAA binding motifs indicated by shaded boxes. Transcription factor binding sites predicted by MATCH to be disrupted are shown in black. Transcription factor binding sites predicted by PROMO to be disrupted are shown in blue.

### 3.7. Transient transfection analysis of Aicda R2-1 mutations

Previous studies have established that mutating the fourth nucleotide within the PU.1 motif from GGAA -> GGA<u>C</u> is sufficient to abrogate its interaction with DNA (Xu et al., 2012). To determine how this mutation would influence the transcriptional repression function of the R2-1 region, site-directed mutagenesis was used to change the GGAA sequence of the R2-1 region to GGA<u>C</u> within the pGL3-R2-1-EiH luciferase reporter plasmid (**Fig. 7A)**. In addition, a mutant was constructed with a 4 bp deletion of the GGAA PU.1 core motif (**Fig. 7A**). Transient transfection into 38B9 cells followed by dual luciferase assays showed that either point mutation or deletion of the PU.1 binding site resulted in a significant increase in luciferase expression compared to the unmutated R1 plasmid (**Fig. 7B, 7C**). This alleviation in repression by mutating the PU.1 binding motif within *Aicda* R2-1 suggests that this transcription factor plays a role in regulating the repressive capabilities of this regulatory element.

**Figure 7.**
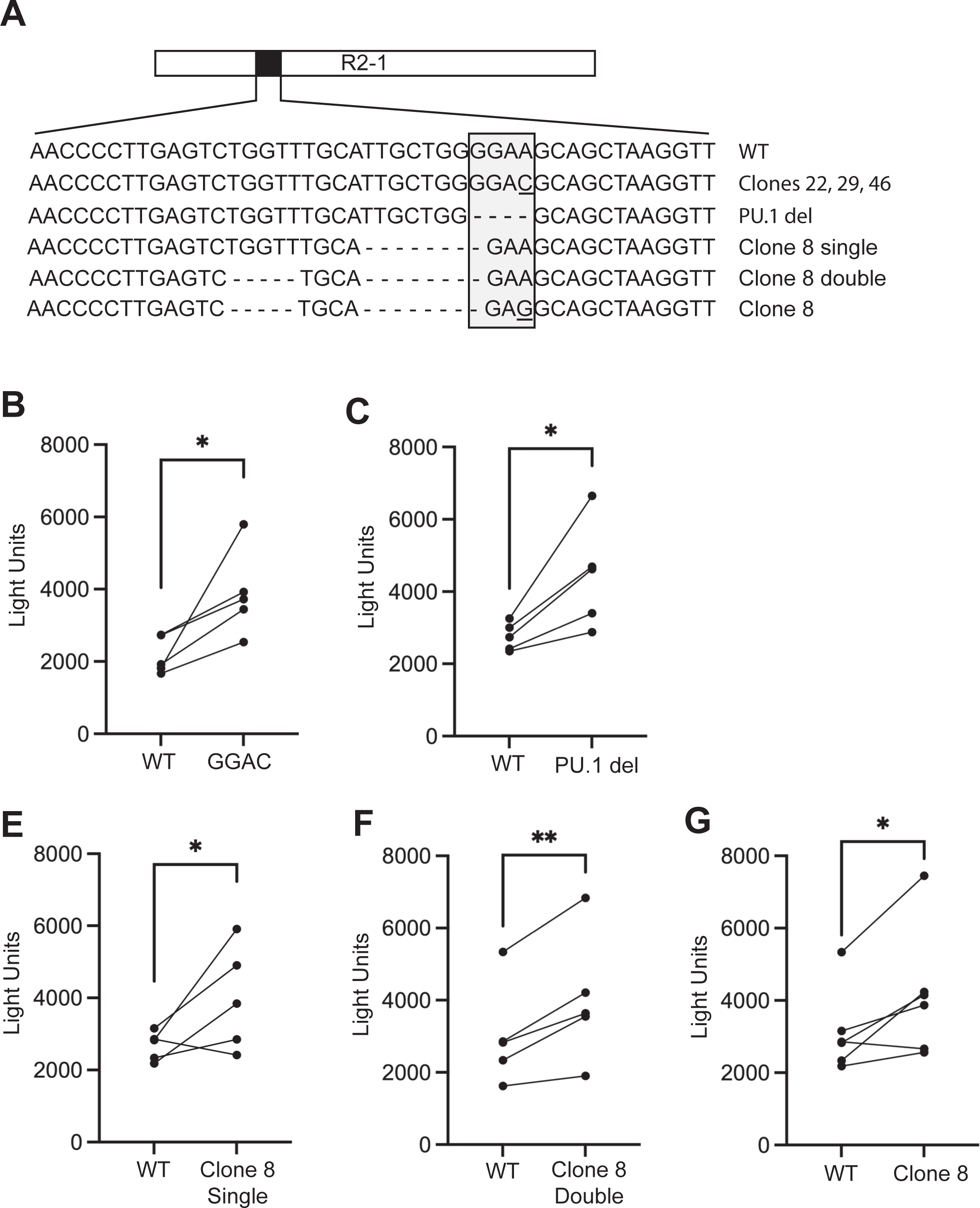
Mutation of the PU.1 interaction site relieves repressive activity of *Aicda* R2-1 in luciferase reporter assays. A) DNA sequences of site-directed mutants of R2-1 in the pGL3-promoter-EiH-R2-1 luciferase reporter plasmid. Box indicates the core GGAA binding motif for PU.1. B-G) PU.1 binding site or clone 8 deletion mutations relieve repression of R2-1. Five independent biological replicate experiments were performed to compare the activity of the wild type with the indicated mutants of the pGL3-promoter-EiH-R2-1 luciferase reporter plasmid. *p<0.05, **p<0.01 by paired t-test. Paired biological replicate experiments are connected by lines.

Next, site-directed mutagenesis was used to introduce the clone 8 mutation into the pGL3-R2-1-EiH reporter plasmid. Three plasmids were generated: the clone 8 single mutation with a 8-bp deletion, the clone 8 double mutation with the 5 bp and 8 bp deletions, and the clone 8 mutation with the 5 bp and 8 bp deletions combined with the A to G point mutation within the PU.1 motif (**Fig. 7A**). Following transient transfection and dual luciferase assay, the plasmid containing the single downstream deletion (pGL3-single) to the R1 region showed an alleviation in repression of the R2-1 region (**Fig. 7E).** Deleting both segments in the pGL3-double deletion plasmid had strong alleviation in luciferase expression relative to the R2-1-containing plasmid, while the clone-8 plasmid had a similar effect as the single deletion (**Fig. 7F, 7G).** This result suggested that the transcription factors predicted to bind within the downstream deletion sequence of clone 8 harbor transcriptional repression capabilities, such that removing these sequences resulted in the upregulation of luciferase expression.

## 4. Discussion

In this study, we found that PU.1 functions as a regulator of *Aicda* transcription in developing B cells. Deletion of *Spi1* encoding PU.1 in cultured leukemia cells or pre-B cells led to up-regulated *Aicda* mRNA transcript levels. Heterozygous CRISPR-Cas9 mutation of the predicted R2-1 regulatory region containing a PU.1 binding site led to upregulated *Aicda* mRNA transcript levels in response to LPS stimulation. R2-1 functioned as a transcriptional repressor in transient transfection analysis followed by luciferase assays. These data suggest that PU.1 is essential for the repressive function of a negative regulator element located in *Aicda* intron 1.

In order to investigate whether *Aicda* might be PU.1-regulated, we determined *Spi1* mRNA transcript levels in “Hardy” stages A-D from developing B cells (**Fig. 1B**). We found dysregulation of *Aicda* mRNA transcripts in fraction B (Hardy et al., 1991). Fraction B contains cells at the pro-B stage of development at which *Cd79a/Mb1* is first expressed, controlling Cre in the Mb1-Cre1′PB mouse model, and is therefore the earliest stage at which PU.1 is deleted (Batista et al., 2017; Hobeika et al., 2006). Leukemia caused by PU.1/Spi-B deletion strongly resembles fraction C large pre-B cells (Batista et al., 2018; Sokalski et al., 2011). We speculate that if driver mutations are induced by dysregulated AID in fraction B, such mutations would be positively selected by proliferation in fraction C. In support of this possibility, AID has been shown to be expressed in human and mouse pro-B cells (Auer et al., 2017; Cantaert et al., 2015). These results suggest that PU.1 is involved in the constraint of *Aicda* transcription in pro-B cells.

We performed experiments to determine if the repression of *Aicda* is mediated by direct PU.1 interaction with region R2-1 within intron 1. Binding of PU.1 to R2-1 was confirmed by ChIP-seq and ChIP-PCR. R2-1 had repressive activity, and point mutation or deletion of the core recognition motif of PU.1 (GGAA) relieved repression in luciferase assays. Heterozygous mutation of R2-1 using CRISPR-Cas9 also de-repressed *Aicda* transcription following LPS stimulation in 38B9 pre-B cells. However, heterozygous point mutations of the PU.1 core binding motif (GGAA -> GGA<u>C</u>) did not affect *Aicda* mRNA transcript levels following LPS stimulation. We speculate that we could not detect dysregulated transcript levels because mutations were heterozygous rather than homozygous. In future studies, homozygous CRISPR-induced mutations will be investigated.

We did not detect any homozygous mutations of R2-1 among the CRISPR clones (**Fig. 4**). Heterozygous mutations are a common result in CRISPR mutagenesis, as the double-stranded DNA breaks created by Cas9 are repaired by non-homologous end joining (NHEJ) rather than homologous recombination, creating heterozygous insertion or deletion mutations (Miyaoka et al., 2016). It is also possible that homozygous mutation of R2-1 of *Aicda* was lethal to 38B9 cells due to upregulation or dysregulation of AID. AID was shown to play a role in B cell tolerance by inducing DNA lesions that led to cell death and removal of autoreactive B cells (Meyers et al., 2011). Overexpression of AID leads to the hyperediting of its target substrate (Martin and Scharff, 2002), causing deregulated somatic hypermutation that could increase the elimination of these cells. Future work will include generating and testing of homozygous R2-1 mutations in cultured cells and *in vivo*.

There were other transcription factors in addition to PU.1, including POU and IKZF family members, predicted to interact with R2-1 identified using PROMO and MATCH software. Prediction of IKZF transcription factor interaction corresponded with our observation that IKZF3 interacts with R2-1 (**Fig. 2**). There are no published reports of POU transcription factors interacting with *Aicda* region 2. Other transcription factors previously shown to interact with region 2 and regulate B cell specific transcription of *Aicda* included Pax-5, E2A, Myb, and E2F (Tran et al., 2010; Zan and Casali, 2013). Taken together, these data suggest that R2-1 is an important regulatory element constraining *Aicda* transcription in developing B cells.

## 5. Conclusions

There is now abundant evidence that off-target effects of AID can induce driver mutations in various types of leukemia (Swaminathan et al., 2015; Zan and Casali, 2013). Mutational signature analysis of pediatric cancers have revealed AID-like signatures of mutagenesis (Swaminathan et al., 2015; Thatikonda et al., 2023). Therefore, it is of paramount importance to understand how DNA lesions affecting transcription factors might regulate *Aicda* transcription during early B cell development. This study shows that a PU.1-interacting regulatory region within intron 1 of *Aicda* is important to constrain transcription. Future work will focus on a comprehensive understanding of how *Aicda* is regulated in a cell type-specific and developmental stage-specific manner.

## Abbreviations

AID: activation-induced cytidine deaminase
ChIP: chromatin immunoprecipitation
pre-B-ALL: precursor B cell acute lymphoblastic leukemia
BCR: B cell receptor
RAG: recombinase activating gene
IL-7: interleukin-7
RT-qPCR: reverse transcriptase quantitative polymerase chain reaction
NCR: negative control region
EiH: immunoglobulin heavy chain intronic enhancer.

## CRediT authorship contribution statement

**Allanna MacKenzie**: Conceptualization, Data Curation, Formal Analysis, Visualization, Methodology, Writing – original draft. **Mia Sams**: Conceptualization, Methodology. **Jane Lin**: Conceptualization, Methodology. **Carolina Reyes Batista**: Conceptualization, Methodology. **Michelle Lim**: Conceptualization, Formal Analysis, Methodology. **Chanpreet Riarh**: Conceptualization, Methodology. **Rodney DeKoter**: Conceptualization, Data Curation, Formal Analysis, Funding Acquisition, Project Administration, Writing – Review and Editing.

## Data Availability

Data will be made available upon request.

## Acknowledgements

We acknowledge Jenn Biltcliffe of the London Regional Genomics Centre for assistance with DNA sequencing. We acknowledge the Centre for Applied Genomics (Toronto, Ontario) for DNA sequencing services. We thank Kristin Chadwick of the London Regional Flow Cytometry Facility for assistance with cell sorting.

## Supporting information

One supplemental Table and one supplemental Figure.

