## Supporting Information for "Negative regulation of Activation-Induced Cytidine Deaminase gene transcription in developing B cells by a PU.1-interacting intronic region"

**Supplemental Information – MacKenzie et al.**

**Supplemental Table 1 – Primer Sequences**

| Primer Name | Forward Primer Sequence | Reverse Primer Sequence |
|---|---|---|
| sgRNA cassette isolation and cloning | 5'- ATG GAA AAA CGC CAG CAA CGC -3' | 5'- CAA TGG GCG GGG GTC GTT GGG -3' |
| Deletion of SOX17 cRNA (Q5 site directed mutagenesis) | 5'- GTT TTA GAG CTA GAA ATA GCA AG –3' | 5'- GGT GTT TCG TCC TTT CCA C -3' |
| *Aicda* R2-1 crRNA insertion (Q5 site directed mutagenesis) | 5'- TTT GCA TTG CGT TTT AGA GCT AGA AAT AGC AAG -3' | 5'- CCA GAC TCA AGG TGT TTC GTC CTT TCC AC -3' |
| Deletion of R2-1 256-263 (Q5 site directed mutagenesis) | 5'- GAA GCA GCT AAG GTT CTG -3' | 5'- TGC AAA CCA GAC TCA AGG -3' |
| Deletion of R2-1 247-251 (Q5 site directed mutagenesis) | 5'- TGC AGA AGC AGC TAA GGT -3' | 5'- GAC TCA AGG GGT TAA AAA TAA TC -3' |
| Clone 8 point mutation R2-1 (Q5 site directed mutagenesis) | 5'- AGT CTG CAG AGG CAG CTA AGG -3' | 5'- CAA GGG GTT AAA AAT AAT CCC AG -3' |
| *Aicda* R2-1 clone screening (genomic DNA template, 355 bp product) | 5'- TGA GCT GCT TCT GGG TTT ATA G-3' | 5'- GTA AGG AGG ACT TTG CTA GGT G -3' |
| Gibson assembly sequencing | 5'- CCG TAA CTT GAA AGT ATT TCG -3' | 5'- TAA TCA GTA GCG ATA ATG GTA -3' |
| *Aicda* intron one (negative control) ChIP-qPCR | 5'- TGC TAC TAG GCT GGG ATT ATT T -3' | 5'- AGT GCT CTT AAC CAC TGA GC -3' |
| *Aicda* PU.1 ChIP-qPCR | 5'- AGG GCA GAA GGT TAA AGG TG -3' | 5'- CGC CAG CTG CTG AGA CA -3' |
| *IgH* EiH amplification for cloning | 5'- ACG GAT CCT CTA GAG AGG TCT GGT GGA GCC TGC -3' | 5'- CCG GAT CCT CTA GAT AAT TGC ATT CAT TTA AAA -3' |
| *Aicda* R2-1 amplification for cloning | 5'-TGA GCT GCT TCT GGG TTT ATA G-3' | 5'-GAG TTT AGA AAG GAT GTG TCT CAA ATA G-3' |
| *Aicda* R2-2 amplification for cloning | 5'-GAA CAG CCA ATA GCG ACA TAG A-3' | 5'-GCC TTA GAA GCA CGG GAA TTA-3' |
| *Tbp* RT-qPCR | 5'-ACC GTG AAT CTT GGC TGT AAA C-3' | 5'-GCA GCA AAT CGC TTG GGA TTA-3' |
| *Aicda* (exon 2-3) RT-qPCR | 5'-CTC CTG CTC ACT GGA CTT C-3' | 5'-GTC TGA GAT GTA GCG TAG GAA C-3' |

**Supplemental Figure 1. Allele-specific sequencing of 38B9 clone 46 and clone 8.** PCR was used to amplify *Aicda* R2-1 products from 38B9 clones 46 and 8, followed by cloning of PCR products and sequencing of individual clones. A) Sequencing of clone 46. 3 clones had a single point mutation and 2 clones were wild type. B) Sequencing of clone 8. 7 clones had the mutation(s) shown on the top and 7 clones had the 2 nt deletion mutation shown on the bottom.
